# *Irx3/5* define the cochlear sensory domain and regulate vestibular and cochlear sensory patterning in the mammalian inner ear

**DOI:** 10.1101/2024.10.24.620152

**Authors:** Yuchen Liu, Tianli Qin, Xin Weng, Bernice Leung, Karl Kam Hei So, Boshi Wang, Wanying Feng, Alexander Marsolais, Sheena Josselyn, Pingbo Huang, Bernd Fritzsch, Chi-Chung Hui, Mai Har Sham

**Author notes:** Y.L., T.Q., X.W. contributed equally to this work.

## Abstract

The mammalian inner ear houses the vestibular and cochlear sensory organs dedicated to sensing balance and sound, respectively. These distinct sensory organs arise from a common prosensory region, but the mechanisms underlying their divergence remain elusive. Here, we showed that two evolutionarily conserved homeobox genes, *Irx3* and *Irx5*, are required for the patterning and segregation of the saccular and cochlear sensory domains, as well as for the formation of auditory sensory cells. *Irx3/5* were highly expressed in the cochlea, their deletion resulted in a significantly shortened cochlea with a loss of the ductus reuniens that bridged the vestibule and cochlea. Remarkably, ectopic vestibular hair cells replaced the cochlear non-sensory structure, the Greater Epithelial Ridge. Moreover, most auditory sensory cells in the cochlea were transformed into hair cells of vestibular identity, with only a residual organ of Corti remaining in the mid-apical region of *Irx3/5* double knockout mice. Conditional temporal knockouts further revealed that *Irx3/5* are essential for controlling cochlear sensory domain formation before embryonic day 14. Our findings demonstrate that Irx3/5 regulate the patterning of vestibular and cochlear sensory cells, providing insights into the separation of vestibular and cochlear sensory organs during mammalian inner ear development.

## Main

The mammalian inner ear is a highly intricate structure containing six separate sensory organs responsible for balance and hearing, each of which contains sensory hair cells^1^. Five vestibular sensory organs, including saccular and utricular macula, and three canal cristae (anterior, posterior, horizontal) sense balance and movement. The cochlea in the ventral part of the inner ear is dedicated for sound perception^1^. Nevertheless, the mechanisms of formation of these distinct sensory patches in the inner ear remain elusive.

*Irx* genes encode evolutionarily conserved TALE homeodomain transcription factors which play important roles during embryogenesis. In mammals, *Irx* genes are grouped into two clusters: *IrxA* cluster contains *Irx1, Irx2* and *Irx4*; *IrxB* cluster includes *Irx3, Irx5* and *Irx6* ^2^. *Irx3* and *Irx5* genes are essential in developmental processes of various systems, including bone, heart, limb, hypothalamus, etc^2-6^. Mutations in *IRX5* gene lead to Hamamy syndrome, a rare developmental disease characterized by craniofacial malformations, congenital heart defects, skeletal anomalies, and sensorineural hearing loss^3^. Interestingly, it has been demonstrated that *Irx3* and *Irx5* are expressed in the developing chick inner ear at multiple stages^7^. However, the function of *Irx3/5* genes in mammalian cochlear development remain unknown.

### Irx3/5 deficiency leads to abnormal inner ear morphogenesis and formation of ectopic HCs of vestibular feature in the cochlea

To understand the roles of *Irx3* and *Irx5* in the mammalian inner ear, we first examined the expression patterns of both genes using *Irx3*^*LacZ/+*^ and *Irx5*^*EGFP/+*^ transgenic knock-in reporter mouse mutants^4^. We observed that both *Irx3* and *Irx5* were broadly expressed in the cochlear epithelium and the surrounding mesenchyme at E14.5 (Supplementary Fig. S1a-b). Consistent with the hearing impairment manifested in human patients with *IRX5* mutations^3^, *Irx3*^*flox*^ *Irx5*^*EGFP*^*/Irx3*^*flox*^ *Irx5*^*EGFP*^ (designated as *Irx5*^*-/-*^) mice displayed increased auditory brainstem response (ABR) thresholds at low frequencies (Fig. 1j,k). Moreover, ABR thresholds assessment in *Irx3*^*LacZ/LacZ*^ (designated as *Irx3*^*-/-*^) mice also showed defective hearing functions, with elevated ABR thresholds at frequencies around 16-20kHz (Fig. 1i,k).

**Fig 1.**
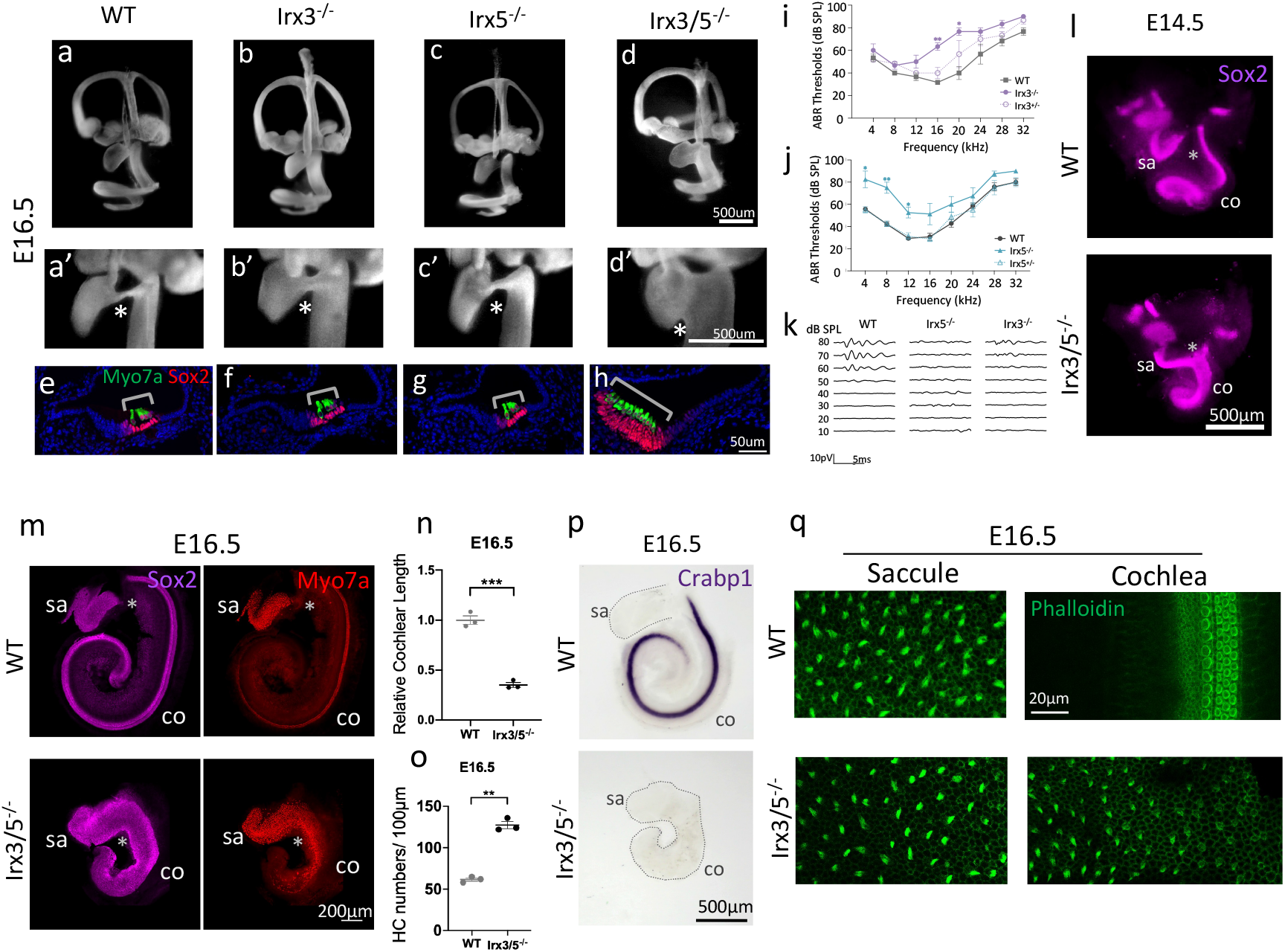
Removal of *Irx3/5* leads to defective inner ear morphogenesis and supernumerary vestibular HC formation in the cochlea. a-d’, Paint-fill analysis of E16.5 WT, *Irx3*^*-/-*^, *Irx5*^*-/-*^ and *Irx3/5*^*-/-*^ inner ears (n=3 for all samples). Cochlear duct was shortened and ductus reuniens was lost (indicated by asterisk) in *Irx3/5*^*-/-*^, while single *Irx3* or *Irx5* mutants were relatively normal. Zoom in view of the ductus reuniens in a-d (a’, b’, c’, d’). e-h, Myo7a and Sox2 immunostaining on E16.5 WT, *Irx3*^*-/-*^, *Irx5*^*-/-*^ and *Irx3/5*^*-/-*^ cochlear sections (n>3 for all samples). Ectopic hair cells were observed and the Sox2^+^ sensory domain was expanded in *Irx3/5*^*-/-*^ cochlea. Formation of the organ of Corti was relatively normal in *Irx3*^*-/-*^ and *Irx5*^*-/-*^ cochlea. i, ABR thresholds for 2-month-old *Irx3*^*-/-*^ mice (n=3), together with heterozygous (n=3) and WT littermates (n=3). Mean±SEM. * p ≤ 0.05, ** p ≤ 0.01, unpaired two-tailed t tests with Welch’s correction. Data suggested hearing impairment of the *Irx3*^*-/-*^ mice under medial frequency (16-20kHz) stimuli. j, ABR thresholds for 2-month-old *Irx5*^*-/-*^ adult mice (n=4) and their littermates, *Irx5*^*+/-*^ (n=4) and WT mice (n=7). Mean±SEM. * p ≤ 0.05, ** p ≤ 0.01, unpaired two-tailed t tests with Welch’s correction. *Irx5*^*-/-*^ mice displayed significantly elevated hearing thresholds under low sound frequency stimuli. k, Representative ABR traces elicited by click stimuli indicate hearing impairments of 2-month-old *Irx3*^*-/-*^ and *Irx5*^*-/-*^ mice. l, Whole mount Sox2 immunostaining of E14.5 WT and *Irx3/5*^*-/-*^ inner ears (n=3 for all samples). All six sensory patches were separated from each other in the WT. However, saccular and cochlear sensory domains were merged in the *Irx3/5*^*-/-*^ inner ear (highlighted by white asterisk). m, Whole mount Sox2 and Myo7a immunostaining in saccule and cochlea of E16.5 WT and *Irx3/5*^*-/-*^ (n=3 for all samples). Sox2^+^ sensory domains and Myo7a^+^ hair cells were well segregated between saccule and cochlea in the WT. Nevertheless, saccular and cochlear Sox2^+^ sensory domains were fused together and Myo7a^+^ hair cells exhibited continuous pattern in *Irx3/5*^*-/-*^. Myo7a^+^ hair cells in *Irx3/5*^*-/-*^ cochlea were located along the medial edge. n, Quantification of E16.5 *Irx3/5*^*-/-*^ cochlear length relative to WT (n=3 for all samples). mean ± SEM. *** p ≤ 0.001, unpaired two-tailed t tests with Welch’s correction. o, Quantification of hair cell numbers in E16.5 WT and *Irx3/5*^*-/-*^ cochlea (n=3 for all samples). mean ± SEM. ** p ≤ 0.01, unpaired two-tailed t tests with Welch’s correction. p, Whole mount *in situ* hybridization of *Crabp1*, a marker for GER region, in E16.5 WT and *Irx3/5*^*-/-*^ cochlea (N=3 for all samples). *Crabp1* expression was absent in the *Irx3/5*^*-/-*^ cochlea. q, Whole mount Phalloidin immunostaining of hair cell stereocilia bundles in E16.5 WT and *Irx3/5*^*-/-*^ cochlear basal region and saccule. Stereocilia bundles of hair cells in *Irx3/5*^*-/-*^ cochlea were long and wispy, resembling that of vestibular hair cells (N=3 for all samples).

To explore the functions of *Irx3* and *Irx5* during inner ear development, we analyzed the morphology of *Irx3*^*-/-*^, *Irx5*^*-/-*^ and *Irx3*^*-*^ *Irx5*^*EGFP*^/*Irx3*^*-*^ *Irx5*^*EGFP*^ (designated as *Irx3/5*^*-/-*^) mutant inner ears. Paint-fill analyses showed that while the gross structures and morphologies of *Irx3*^*-/-*^ and *Irx5*^*-/-*^ inner ear were comparable to that of control (Fig. 1a-c’), cochlear duct was shortened and the ductus reuniens, a fine non-sensory structure that connects the saccule and cochlea, was absent in *Irx3/5*^*-/-*^ (Fig. 1a,a’,d,d’). Immunostaining of Myo7a and Sox2 showed an expansion of sensory region to the medial edge of the floor epithelium and formation of ectopic HCs in E16.5 *Irx3/5*^*-/-*^ cochlea (Fig. 1e,h). In contrast, the formation of the organ of Corti (oC) was largely normal in the cochlea of *Irx3*^*-/-*^ or *Irx5*^*-/-*^ single mutants (Fig. 1f,g). Collectively, these results showed that *Irx3* and *Irx5* compensate each other’s function in mammalian inner ear development and removal of both *Irx3* and *Irx5* leads to defective inner ear morphogenesis and abnormal HC development in the cochlea.

Next, we sought to understand the underlying reasons of ectopic HC formation and expanded sensory region in *Irx3/5*^*-/-*^ cochlea. At E14.5, while all six Sox2^+^ sensory domains in the WT inner ear were separated from each other, saccular sensory domain remained fused with the cochlear sensory domain in *Irx3/5*^*-/-*^ (Fig. 1l). At E16.5, a clear gap was observed between the Sox2^+^ sensory domains in saccule and cochlea, although they were still physically connected in the WT; Myo7a+ HCs were formed within the Sox2+ sensory regions and cochlear HCs differentiated from base to apex (Fig. 1m). Strikingly, in *Irx3/5*^*-/-*^, the cochlear duct was significantly shortened, saccular and cochlear Sox2^+^ sensory regions were fused together without any gap, and ectopic HCs were formed in the medial portion of the cochlear floor epithelium (Fig. 1m-o). The ectopic HCs located medially to the presumptive oC in *Irx3/5*^*-/-*^ cochlea occupied the normal cochlear GER. *Crabp1*, a differentiating GER marker^8^, was expressed in the GER of E16.5 WT cochlea from the base to apex (except for the apical end). However, its expression was completely absent in *Irx3/5*^*-/-*^ cochlea (Fig. 1p), suggesting the loss of GER identity which was replaced by sensory cells. Interestingly, stereocilia bundles of the HCs in *Irx3/5*^*-/-*^ cochlea were long and wispy, resembling that of vestibular HCs^9^ (Fig. 1q). These results demonstrated a failure of segregation of cochlear and saccular sensory domains, and the transformation of non-sensory cells in the cochlear GER region into sensory cells with vestibular features when both *Irx3* and *Irx5* functions were lost.

### Cochlear and saccular sensory domains gradually separate from each other from E12.5 to E14.5 and requires *Irx3/5*

To understand the formation and segregation of saccular and cochlear sensory regions during inner ear development, we characterized Sox2 and Myo7a expression at different developmental stages using whole mount immunostaining. At E12.5, Sox2 expression showed that these two sensory domains are fused as one sensory patch in both WT and *Irx3/5*^*-/-*^ inner ears (Supplementary Fig. S2a-b’’). This merged sensory region gradually segregates into two discrete parts located in the saccule and cochlea from E12.5 to E14.5 in the WT (Fig. 2a,a’, Supplementary Fig. S2a-a’’, c-c’’). However, this segregation failed to occur in *Irx3/5*^*-/-*^ mutants (Fig. 2a,a’, Supplementary Fig. S2b-b’’, d-d’’). While saccular and cochlear sensory organs were fused at E12.5, HCs were only found in the saccule, which is consistent with the notion that vestibular HCs differentiate earlier than cochlear HCs^10,11^ (Supplementary Fig. S4a-a’’). From E12.5 to E14.5, Myo7a^+^ saccular HCs were developing inside the saccule of the WT (Fig. 2a,a’, Supplementary Fig. S2a-a’’, c-c’’). On the contrary, in *Irx3/5*^*-/-*^, Myo7a^+^ HCs gradually reach to the cochlear mid-apical region along the medial edge of the Sox2^+^ sensory domain from E12.5 to E14.5, forming a continuous band of HCs connected to the saccular sensory patch (Fig. 2a,a’, Supplementary Fig. S2b-b’’, d-d’’). These results indicated an early requirement of *Irx3* and *Irx5* in regulating inner ear sensory patterning.

**Fig 2.**
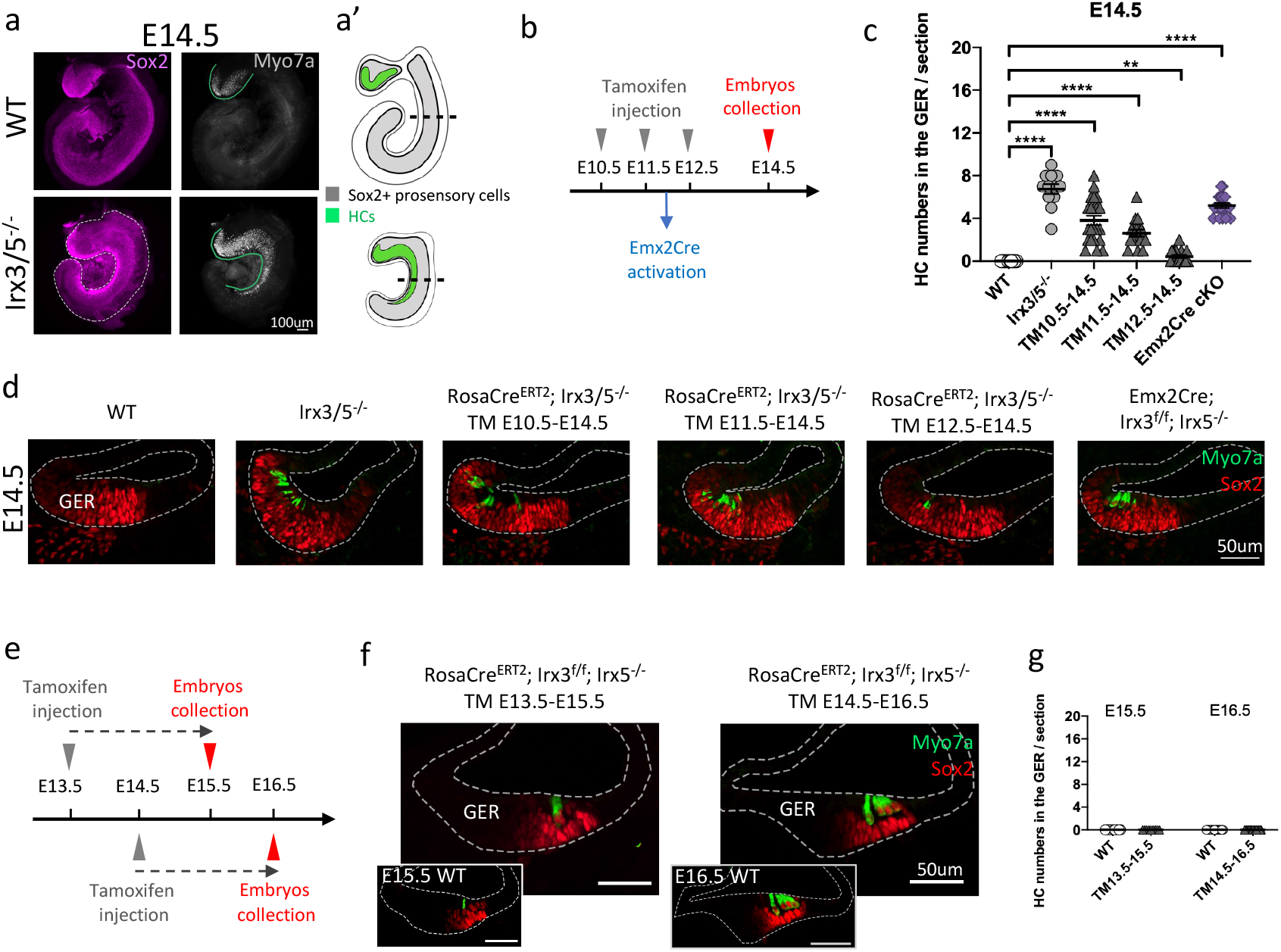
Removal of *Irx3/5* leads to defective inner ear morphogenesis and ectopic vestibular HC formation in the cochlea. a, Whole mount Sox2 and Myo7a immunostaining in saccule and cochlea of E14.5 WT and *Irx3/5*^*-/-*^ (N=3 for all samples). While saccule and cochlear sensory patches were separated in WT, they were merged in *Irx3/5*^*-/-*^. Ectopic hair cells were found along the medial portion of the cochlear floor epithelium and related to the saccular hair cells in *Irx3/5*^*-/-*^. A schematic diagram was shown in a’. b, Summary of tamoxifen injection time points and *Emx2Cre* activation time point to generate mutants for experiments shown in d. Embryos were collected at E14.5. c, Quantification of ectopic hair cell numbers in the GER region from the experiments shown in d. mean ± SEM. ** p ≤ 0.01, **** p ≤ 0.0001, unpaired two-tailed t tests with Welch’s correction. d, Sox2 and Myo7a immunostaining on the sections of cochlea from E14.5 WT, *Irx3/5*^*-/-*^, *Emx2Cre*;*Irx3*^*flox*^*Irx5*^*EGFP*^*/Irx3*^*flox*^*Irx5*^*EGFP*^ and various *RosaCre*^*ERT2*^;*Irx3*^*flox*^*Irx5*^*EGFP*^*/ Irx3*^*flox*^*Irx5*^*EGFP*^ with tamoxifen injected at different time point (N≥3 for all samples). e, Summary of tamoxifen injection time and embryo harvest time for experiments in f. f, Sox2 and Myo7a immunostaining on the sections of E15.5 *RosaCre*^*ERT2*^;*Irx3*^*flox*^*Irx5*^*EGFP*^*/Irx3*^*flox*^*Irx5*^*EGFP*^ cochlea with tamoxifen injected at E13.5 and E16.5 *RosaCre*^*ERT2*^;*Irx3*^*flox*^*Irx5*^*EGFP*^*/Irx3*^*flox*^*Irx5*^*EGFP*^ cochlea with tamoxifen injected at E14.5, together with staining of the WT cochlea (N=3 for all samples). g, Quantification of ectopic hair cell numbers in the GER region from the experiments shown in f. Note the increase of a single row of IHC (E15.5) to four rows of IHC and OHCs (E16.5) within the oC, but no HCs were observed in the GER.

Based on the observation of these ectopic HC formation in *Irx3/5*^*-/-*^ cochlea at E14.5, we asked whether *Irx3/5* were continuously required and whether late deletion of *Irx3/5* will also lead to the non-sensory to sensory cell fate conversion in the cochlea. To address the temporal requirement of *Irx3/5*, we generated *RosaCre*^*ERT2*^;*Irx3*^*flox*^ *Irx5*^*EGFP*^*/Irx3*^*flox*^ *Irx5*^*EGFP*^ mutants to delete *Irx3* at different time points in the *Irx5* null background upon tamoxifen injection (Fig. 2b). At E14.5, Sox2 was expressed from the middle to medial side of the cochlear floor epithelium and no Myo7a^+^ HCs were found in the WT cochlea (Fig. 2d). However, HCs could be detected in the medial edge of *Irx3/5*^*-/-*^ cochlea floor epithelium (Fig. 2d), consistent with the whole mount results (Fig. 2a,a’). Similar to *Irx3/5*^*-/-*^, HCs were found in the medial edge of the cochlear floor epithelium from *RosaCre*^*ERT2*^;*Irx3*^*flox*^ *Irx5*^*EGFP*^*/Irx3*^*flox*^ *Irx5*^*EGFP*^ mutants with tamoxifen injected at E10.5, E11.5 and E12.5 (Fig. 2c,d). Consistently, *Emx2Cre*;*Irx3*^*flox*^ *Irx5*^*EGFP*^*/Irx3*^*flox*^ *Irx5*^*EGFP*^ mutant with Cre activation started at around E12 also recapitulate *Irx3/5*^*-/-*^ cochlea phenotype at E14.5 (Fig. 2c,d). Interestingly, when tamoxifen was injected at E13.5 (embryos harvested at E15.5) or E14.5 (embryos harvested at E16.5) (Fig. 2e), HC and oC developed normally as they located in the middle of the cochlear floor without any ectopic HCs forming on the medial edge of the cochlear floor (Fig. 2f,g). Together, these data demonstrated that the requirement of *Irx3/5* function before E14 for inner ear sensory patterning and that late removal of *Irx3/5* was unable to induce non-sensory to sensory identity switch in the cochlea.

### Single cell transcriptome and gene expression analysis reveal abnormal growth of vestibular HCs in *Irx3/5*^x*-/-*^ cochlea

To investigate the roles of *Irx3/5* in specific cell types during inner ear sensory development, we performed single-cell RNA-sequencing (scRNA-seq) analysis of both saccule and cochlea from E14 WT and *Irx3/5*^*-/-*^ embryos (Fig. 3a). We profiled 11590 cells (6787 cells from WT and 4803 cells from *Irx3/5*^*-/-*^ mutant) and annotated different groups of cells using specific markers (Supplementary Fig. S3a-b). Sox2^+^/Jag1^+^ (pro-)sensory cells were then *in silico* isolated from the dataset (Fig. 3b,c). These cells could be identified as saccular sensory cells or cochlear sensory cells based on distinct gene expression profiles (Fig. 3e). We observed an increase of the proportion of the saccular sensory cells in *Irx3/5*^*-/-*^ mutants (Fig. 3d). Interestingly, *Irx3* and *Irx5* are two genes highly enriched in the cochlea (Fig. 3e). Consistently, EGFP and LacZ expression also confirmed that both *Irx3* and *Irx5* were predominantly expressed in the cochlea at E13.5 (Fig. 3f). In addition, we also identified several genes which are abundantly expressed in the saccule or cochlea (Fig. 3e). To examine the identity of sensory cells in *Irx3/5*^*-/-*^ cochlea, we performed in situ hybridization using a specific molecular marker *Tnfaip2*, a gene found to be expressed in the saccule but not in the cochlea. Single cell analysis^12^ of postnatal inner ear also confirmed that *Tnfaip2* was specifically expressed in the vestibular supporting cells (Supplementary Fig. S4). Due to the embryonic lethality of *Irx3/5*^*-/-*^ mutant embryos, we analyzed E17.5 mutants and WT. In the WT, *Tnfaip2* was expressed broadly within the saccule and no expression was detected in the cochlea (Fig. 3g). Remarkably, in *Irx3/5*^*-/-*^ mutants, we detected an ectopic expression of *Tnfaip2* in the cochlea (Fig. 3g). Co-stain of Myo7a showed that HCs in the cochlea overlap with the *Tnfaip2*^+^ region, demonstrating a conversion of cochlear sensory organ to the vestibular sensory organ when both *Irx3* and *Irx5* were inactivated (Fig. 3g).

**Fig 3.**
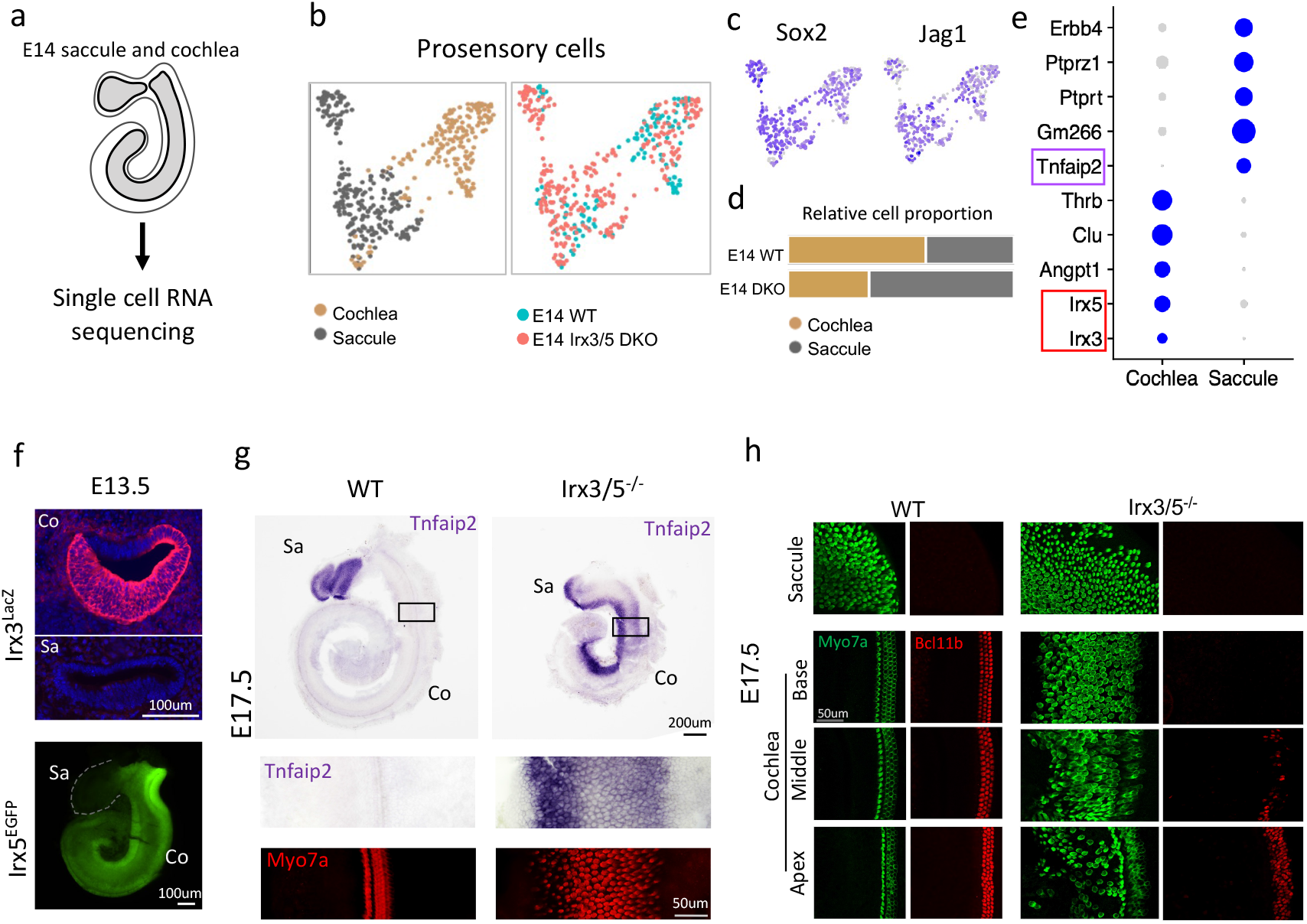
scRNA-seq and gene expression revealed abnormal vestibular hair cell development in the *Irx3/5*^*-/-*^ cochlea. a, Cochlea and saccule from E14 WT and *Irx3/5*^*-/-*^ inner ears were dissected out and processed to single cell RNA sequencing (scRNA-seq). b, UMAP plot of *in silico* isolated prosensory cells from scRNA-seq dataset. Identities and genotypes of cells are indicated. c, Featureplots show *Sox2* and *Jag1* expression in prosensory cells in b. d, Relative proportion of cochlear prosensory and saccular prosensory cells in WT and *Irx3/5*^*-/-*^ mutant. e, Dotplot of differentially expressed genes in saccule and cochlea. Purple circle highlights saccular specific gene *Tnfaip2*, which is also validated in g. Red circle highlights both *Irx3* and *Irx5* are cochlear specific genes, which are also validated in f. f, LacZ and EGFP reporters show *Irx3* and *Irx5* are predominantly expressed in the cochlea, but not in the saccule at E13.5. g, Whole mount *in situ* hybridization of *Tnfaip2* counterstained with Myo7a in the cochlea and the saccule from E17.5 WT and *Irx3/5*^*-/-*^. Black box regions in the cochlea are highlighted in the lower panel. While *Tnfaip2* was only expressed in the WT saccule, ectopic expression of *Tnfaip2* was detected in *Irx3/5*^*-/-*^ cochlea (N≥3 for all samples) and colocalize with Myo7a^+^ hair cells. h, Whole mount immunostaining of Bcl11b and Myo7a in the saccule and cochlea from E17.5 WT and *Irx3/5*^*-/-*^ (n≥5 for all samples). Bcl11b marks cochlear outer hair cells. Bcl11b expression was only detected in the mid-apical region of *Irx3/5*^*-/-*^ cochlea.

We further examined whether cochlear HCs could be formed in the mutants. We analyzed the presence of cochlear HCs by examining the expression of Bcl11b, an cochlear outer hair cell (OHC) specific marker^13,14^. At E17.5, while Bcl11b expression was not found in the saccule, its expression was detected specifically in the OHCs throughout the cochlea from base to apex (Fig. 3h). By contrast, in the mutant cochlea, Bcl11b expression was restricted to the lateral part of the Myo7a^+^ HC domain in the mid-apical regions (Fig. 3h). Based on the ectopic *Tnfaip2* expression and the absence of Bcl11b^+^ OHCs, HCs in the basal parts of the mutant cochlea adopted a vestibular HC fate. Interestingly, although residual oC existed in the mutant cochlear apex, ectopic HCs were found in the GER region of the apex. Therefore, as revealed by the loss of *Crabp1* (Fig. 1p), ectopic *Tnfaip2* expression in the medial portion of the floor epithelium from base to apex (Fig. 3g) and ectopic Myo7a^+^ HCs (Fig. 3h), the non-sensory GER along the entire cochlear basal-apical axis of *Irx3/5* mutants were transformed into sensory cells of vestibular features.

## Discussion

Combining transgenic mouse mutants and scRNA-seq analyses, our study uncovered essential roles of *Irx3* and *Irx5* in the segregation of sensory regions in the saccule and cochlea. Saccular and cochlear sensory domains gradually separate from each other from E12.5 to E14.5 and this process requires *Irx3/5*. Removal of both *Irx3* and *Irx5* leads to fusion of saccular and cochlear sensory domains, and conversion of cochlear non-sensory GER region into sensory cells of vestibular identity. Both *Irx3* and *Irx5* are highly expressed in the cochlea which highlight their critical roles in securing cochlear cell fate and ensuring the precise patterning of inner ear sensory organs. Notably, a vestigial oC forms at the cochlear apical region in *Irx3/5* mutants, indicating that other inductive signals or genes could potentially activate the cochlear sensory program.

All sensory patches in the inner ear share the characteristics of expressing Sox2 and Jag1. At E12.5, Sox2^+^ sensory domains in cochlea and saccule are fused together in the WT. However, HC differentiation already started in the saccule but not in the cochlea, indicating there are differential molecular regulations of prosensory differentiation between saccule and cochlea at E12.5. This integrated sensory patch was gradually separated and became two distinct sensory organs at E14.5. The ductus reunien, a fine structure will then form in between the saccule and cochlea. Remarkably, this separation process does not occur in the *Irx3/5* mutant inner ear. Moreover, at E13.5, while saccular HCs are developing inside the chamber of saccule in the WT, we could already detect ectopic HCs along the medial edge of the *Irx3/5* mutant cochlea, connecting the saccular sensory region. This ectopic HC differentiation extended to the mid-apex of the *Irx3/5* mutant cochlea at E14.5. Therefore, *Irx3/5* are not only required for separating the saccular and cochlear sensory organs, but also for preventing cells in the cochlea to adopt vestibular cell fate. As *Irx3* or *Irx5* single mutants do not have significant defects of sensory development, there is evident compensation between these two genes in regulating inner ear sensory patterning.

Using tamoxifen induced timed deletion of *Irx3/5*, we uncovered the temporal requirement of *Irx3* and *Irx5*, specifically within E12.5 and E13.5, in regulating the sensory patterning of saccule and cochlea. Remarkably, when *Irx3/5* are deleted after E13.5, it did not lead to overt defects in the cochlea, indicating cochlear GER cells are already specified at E13.5 and removal of *Irx3/5* afterwards will not change their non-sensory identity.

Lineage tracing experiments showed that the entire non-sensory region in the medial side of the cochlear floor and the sensory oC were derived from Sox2^+^ prosensory cells at E12.5^15,16^. Interestingly, these GER and interdental cells within the cochlear medial floor epithelium down-regulate Sox2 expression during development and become non-sensory structures (e.g. inner sulcus), which are important for hearing functions. Conversion of non-sensory GER cells to sensory HCs with vestibular features in *Irx3/5*^*-/-*^ cochlea revealed that these non-sensory GER cells derived from Sox2 prosensory lineage could be directed to sensory cell fate if inhibitory cues are absent.

In sum, in this study, we discovered novel functions of Irx3/5 in regulating the patterning of sensory domains during mammalian inner ear development, and therefore their essential roles in shaping the cochlear territory. These results highlight an involvement of *Irx3/5* in the gene regulatory modules for forming discrete inner ear sensory organs and facilitate our understanding of the development and evolution of mammalian cochlea.

## Materials and Methods

### Animal

All experimental protocols, including the use of *Irx3*^*LacZ* (4)^, *Irx3*^*flox*^*/Irx5*^*EGFP* (4)^, *Irx3*^*-*^*/5*^*EGFP* (4)^, *Emx2Cre* ^(17)^, *RosaCre*^*ERT2* (18)^ mice, were approved by the Committee on the Use of Live Animals in Teaching and Research of the University of Hong Kong (CULATR 4771-18 and 5025-19) and by the Animal Experimentation Ethics Committee of the Chinese University of Hong Kong (20-185-GRF). The mice were housed in the facilities of the Centre for Comparative Medicine Research (CCMR) at the University of Hong Kong and the Laboratory Animal Services Centre at the Chinese University of Hong Kong. The day when a vaginal plug was observed was considered embryonic day 0.5 (E0.5). The pregnant dams were weighed, and each of them received a single tamoxifen dosage at 0.1 mg/g through an intraperitoneal injection.

### Auditory brainstem response (ABR) test

The ABR measurements were performed on 2-month-old mice following previously published procedures^19^. The distance between the external ear of the animal and the MF1 multifield magnetic speaker [Tucker-Davis Technologies (TDT), Alachua, FL, USA] remained consistent for each test. The recording electrode was subdermally inserted at the vertex, the reference electrode at the pinna, and a ground electrode near the tail. For ABR recordings, we utilized RZ6-based TDT (Tucker-Davis Technologies) system III hardware and BioSigRZ software for stimulus presentation and response averaging.

### Immunostaining

The embryonic heads were fixed in 4% paraformaldehyde (PFA) overnight at 4°C. Samples for whole mount immunostaining were further dissected to expose the inner ear sensory epithelia. For immunostaining on sections, after fixation, the samples were immersed in a 15% sucrose solution overnight at 4°C and then embedded in gelatin. The embedded samples were cryo-sectioned at a thickness of 10μm. To prepare the sections for immunofluorescence staining, they were blocked with 10% horse serum for 1 hour at room temperature. The primary antibodies used in this study were Sox2 (Neuromics, GT15098, 1:500), Myo7a (Proteus, 26-6790, 1:500), EGFP (Rockland, 600-101-215, 1:500), galactosidase (Abcam, ab9361, 1: 500), phalloidin (Alexa Fluor, 1:200), Bcl11b (Abcam, ab18465, 1:100), each diluted in 10% horse serum and incubated overnight at 4°C. After washing with PBS, the sections were incubated with secondary fluorescent antibodies (Invitrogen Alexa Fluor) together with DAPI for 1 hour at room temperature. Finally, the fluorescent images were acquired using an Olympus BX51/BX53 fluorescence microscope or a Nikon Ni-U Eclipse upright microscope or an Olympus FV1200 inverted confocal microscope.

### Paint-fill analysis

Inner ear paint-fill was performed as previously described^20^. Briefly, E16.5 mouse embryos were rinsed with PBS, followed by fixation overnight using Bodian’s fixative. The embryos’ heads were bisected and incubated in ethanol overnight, followed by the clearance with methyl salicylate. To observe the inner ear, a glass capillary needle was used to inject 1% white gloss paint into the membranous labyrinth’s lumen through the cochlear duct and utricle. The images were acquired using the Leica MZ10F fluorescence-stereo microscope.

### In situ hybridization

The fixed and dehydrated embryos were rehydrated by shaking them in a decreasing series of methanol in DEPC PBST for 10 minutes each (75%, 50%, 25%, and 0% methanol). The inner ears of the embryos were then dissected in PBST and treated with proteinase K in PBST for about an hour. After rinsing twice, the samples were post-fixed with 4% PFA with 0.2% glutaraldehyde in PBST for 20 minutes. The samples were then sequentially washed and incubated with hybridization mix containing the DIG-labeled RNA probe. After overnight incubation, the samples were washed multiple times and incubated with AP-anti-DIG antibody (Roche). Finally, the hybridization signal was developed by incubating the samples with BM purple AP substrate in the dark at room temperature. The reaction was stopped by washing with PBST, and the inner ears were photographed using a Nikon Ni-U Eclipse upright microscope or a Zeiss Stemi 508 microscope.

### Probe labeling

Plasmids were linearized by restriction enzyme digestion overnight at 37°C. The digested products were then extracted through electrophoresis in 1% DEPC-SeaPlaque GTG Agarose Gel and equilibrated in 1.5ml of 1X β-agarase buffer for 1h at room temperature. The gel was melted at 70°C for 15min, and cooled down at 42°C for 5min. 1 unit of β-agarase (Lonza) for every 200mg of 1% gel was added to the melted gel, followed by incubation at 42°C for 1h. DNA was precipitated with 5M Ammonium Acetate of the same volume, 3-4 times volume of absolute ethanol, and 3ul glycogen (10ug/ul) at room temperature for 30min, then centrifuged at max speed for 30min. The pellet was washed with 1ml 70% DEPC-ethanol and centrifuged for 10min. This washing step was repeated 3 times. The pellet was then dried and dissolved in 45μl DEPC-H2O. 1ug DNA product was incubated at 37°C for 2h together with Transcription Buffer, Dig Labelling Mix, 1μl RNase Inhibitor and 2μl RNA polymerase for a 20μl reaction system, followed by treatment of 2 units DNase I at 37°C for 15min. 2ul 0.5M EDTA (PH 8.0) was added to stop the reaction. 2.5μl 4M LiCl and 75μl prechilled absolute ethanol were added to the reaction system and incubated at -80°C for 30min or at -20°C for 2h to allow precipitation. After centrifugation at 13000g for 15min at 4°C and air dry, the pellet was dissolved in 100μl DEPC-H2O at 37°C for 30min. *Crabp1* probe was kindly provided by Dr. Pierre Chambon^21^. A 355bp fragment of cDNA coding for *Tnfaip2* gene was cloned using the following primer: F: CACCTGCACCTAGTGAAAGAA; R: CTCCCGTGTTGATGTCCAGT, and inserted into pBluescript II KS(+) to generate *Tnfaip2* probe plasmid.

### Single-cell RNA sequencing and analysis

Four inner ears from two WT embryos and two *Irx3/5*^*-/-*^ mutant embryos (one inner ear from one embryo) were collected for dissection. The ventral parts of the inner ears including saccules and cochlear ducts were dissected in cold DEPC-PBS. Dissected samples were digested in 150µl HBSS with 1mg/mL collagenase/dispaseII for 25min at 37°C at 900rpm. 150µl 20% FBS in HBSS was used to stop reaction. Dissociated cells were then filtered using 40μm strainers, centrifuged at 300g for 5min and resuspended in 10% FBS in HBSS to around 1000 cells/µl. Library preparation was performed by Single Cell Omics Core at the School of Biomedical Sciences, the Chinese University of Hong Kong, following the standard protocol of 10X Chromium Next GEM Single Cell 3’ Reagent Kits v3.1. Sequencing was performed using Illumina NextSeq 2000 System at the Chinese University of Hong Kong.

Sequences were aligned to mm10-2020-A using Cell Ranger 7.1.0 (10X Genomics). Processing and analyses of the Cell Ranger output data was done with Seurat (R version 4.0.2) following the tutorial (https://satijalab.org/seurat/index.html). Genes detected in at least three cells were included in the analysis. Cells with unique feature counts over 10000 or less than 200 were filtered out. Cells with more than 10% mitochondrial genes or with more than 50000 total number of RNA molecules were excluded for downstream steps. After filtering, 11590 cells were for downstream analyses.

## Acknowledgements

We thank Shelia Tsang, Elaine Wong and MHS lab members for their suggestions and help throughout this project. This work was supported by the Hong Kong Research Grant Council (RGC GRF 17115520 and 776312). C.C.H.’s research was supported by Canadian Institutes of Health Research and the Canada Research Chairs program. B.F.’s research was supported by National Institute of Aging (AG060504).

---

**Fig S1.**
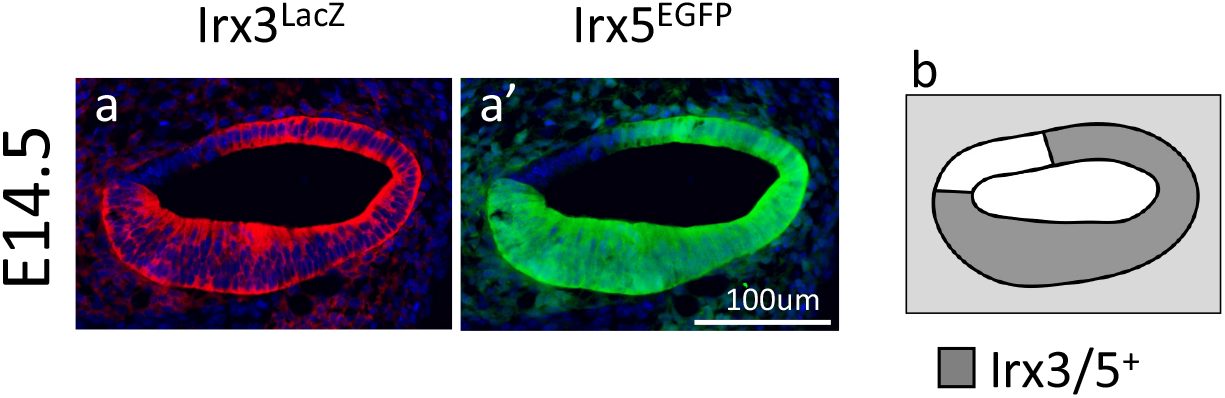
Expression of *Irx3* and *Irx5* in E14.5 cochlea. a, LacZ expression showing that *Irx3* was broadly expressed in the cochlear epithelium and surrounding mesenchyme of E14.5 *Irx3*^*LacZ*^*Irx5*^*+*^*/Irx3*^*+*^*Irx5*^*EGFP*^ embryos (N=3). a’, Costain of EGFP in a showing that *Irx5* was also broadly expressed in the cochlear epithelium and surrounding mesenchyme. b, Schematic diagram of *Irx3* and *Irx5* expression in E14.5 cochlea. These two genes share similar expression patterns.

**Fig S2.**
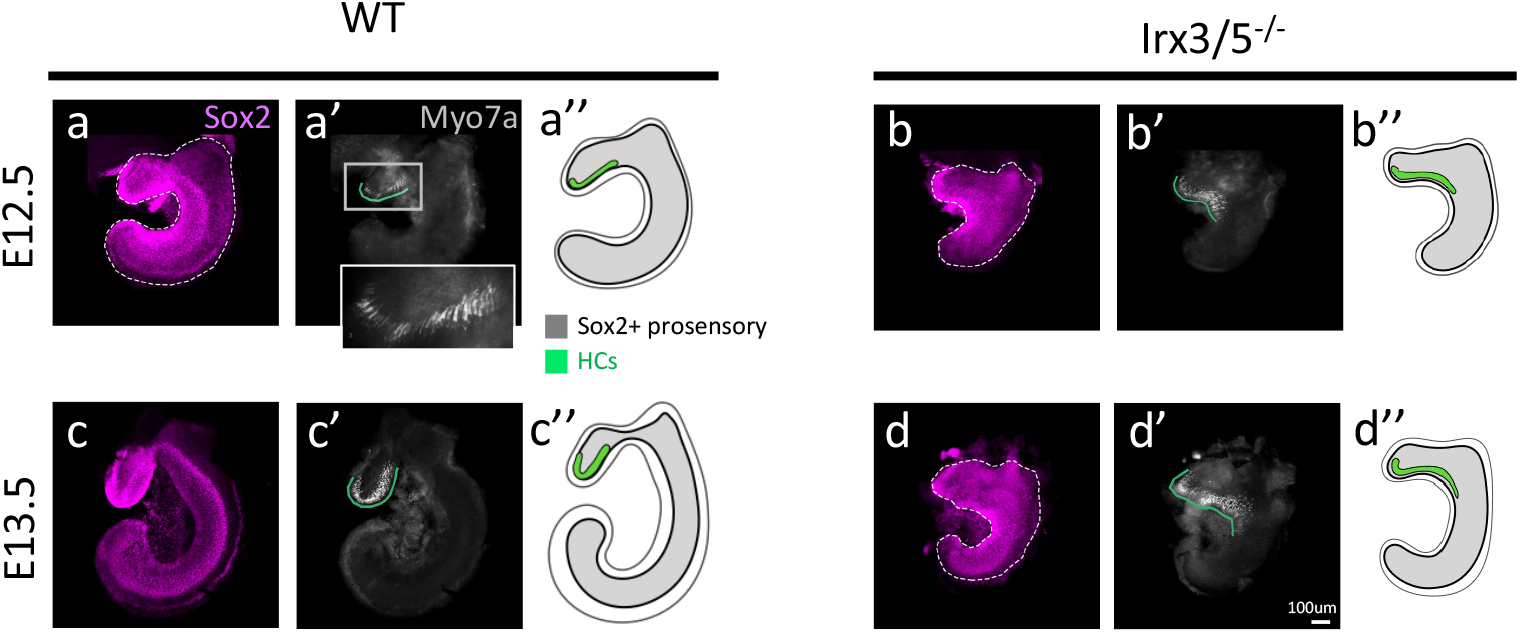
Cochlear and saccular sensory domains gradually separate from each other from E12.5 to E14.5 and requires *Irx3/5*. a-d’’, Whole mount Sox2 and Myo7a immunostaining on saccule and cochlea of E12.5, E13.5 WT and *Irx3/5*^*-/-*^ (N=3 for all samples). Cochlear and saccular sensory domains were fused at E12.5, and they gradually separate from each other from E12.5 to E14.5 (See also Fig. 2a). This process was regulated by *Irx3/5*. Schematic diagram of Sox2 and Myo7a expression in WT and mutants (a’’,b’’,c’’,d’’).

**Fig S3.**
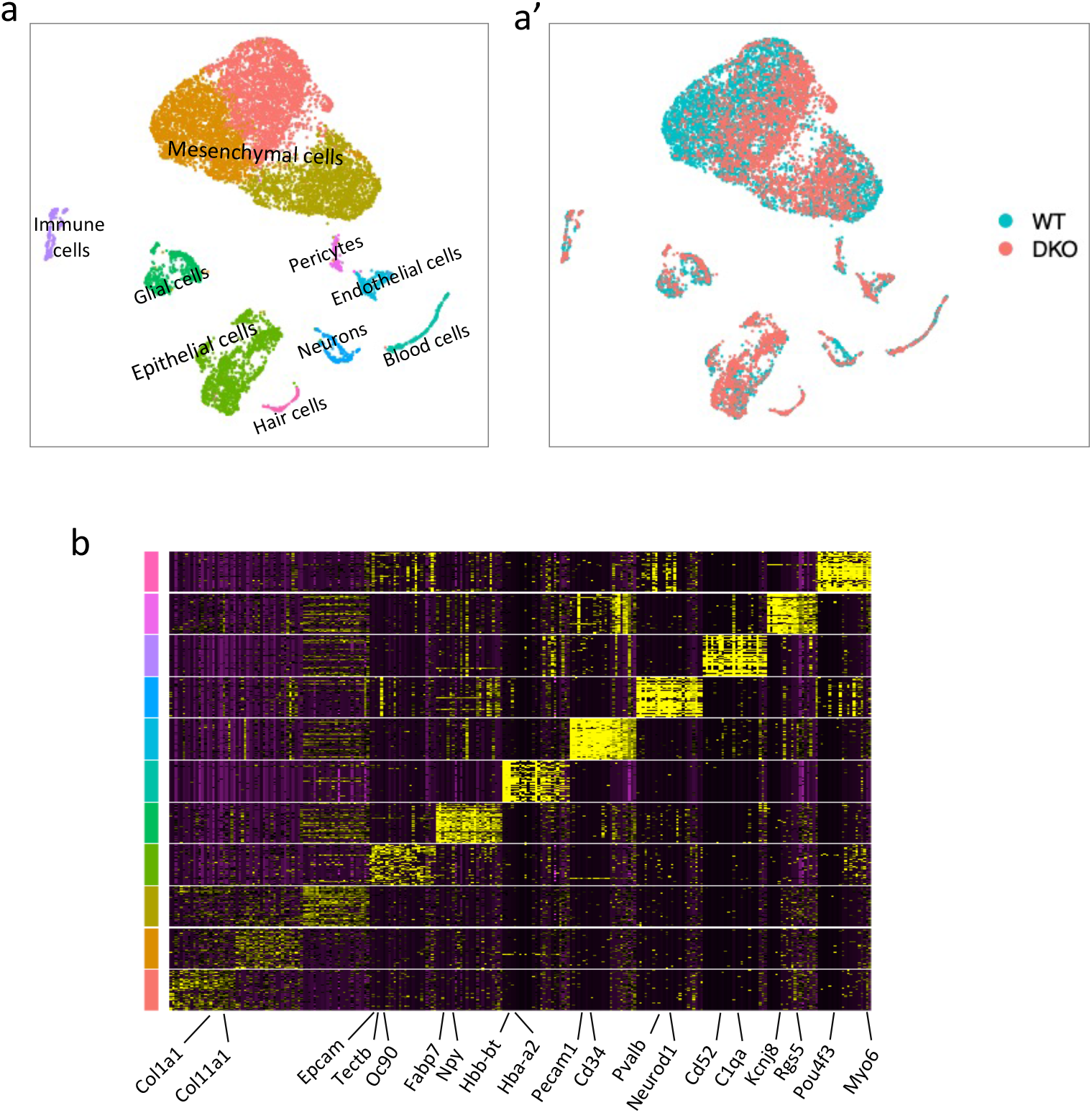
Characterization of transcriptomic profiles of cochlea and saccule from E14 WT and *Irx3/5*^*-/-*^. a, UMAP plot of cochlear and saccular cells profiled by single cell RNA sequencing. Identities of cell clusters were annotated in the plot. a’, Genotypes of each cell in a. b, Heatmap of top 25 markers of each cluster in a.

**Fig S4.**
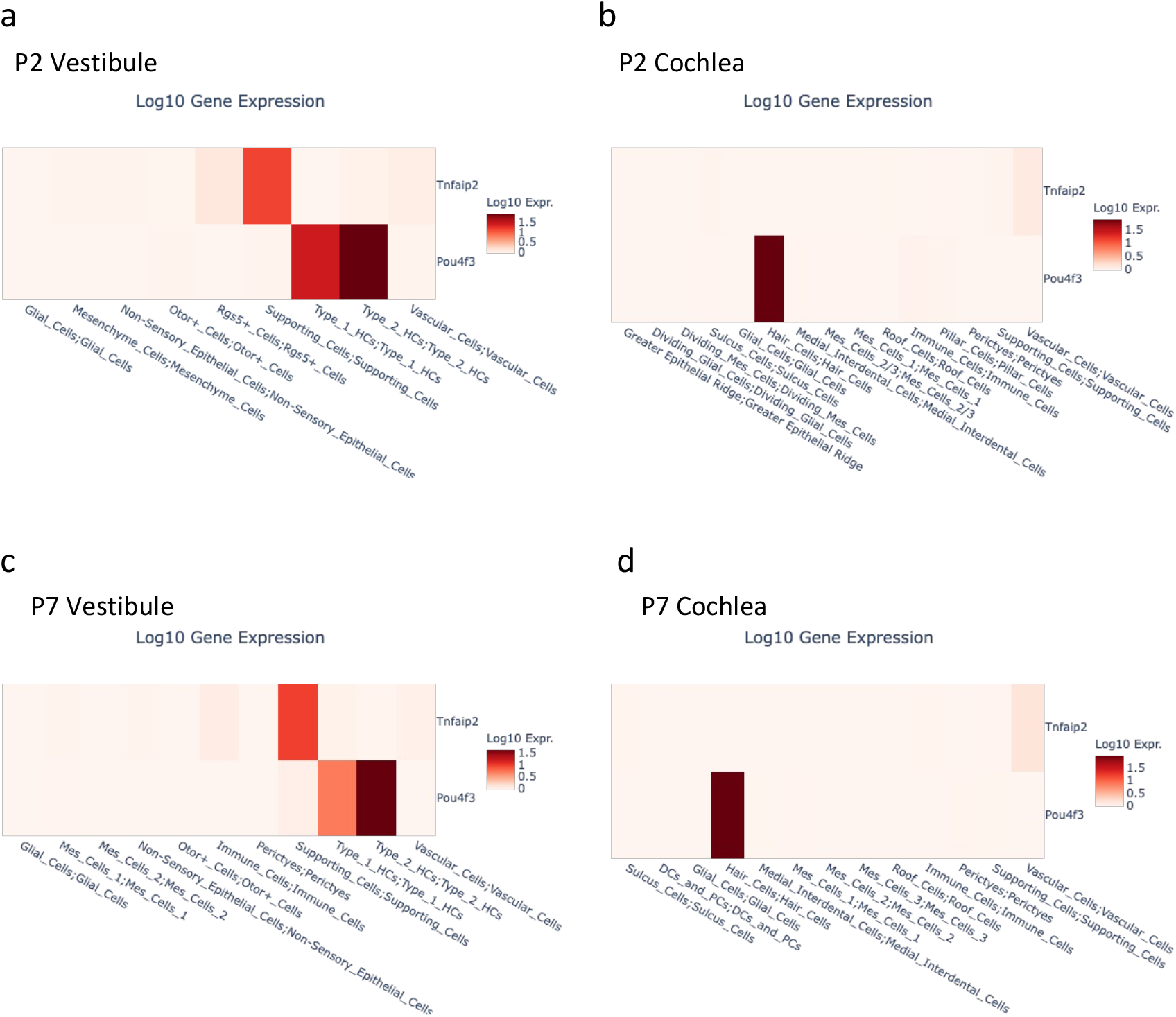
*Tnfaip2* was specifically expressed in the vestibular supporting cells. a-d, Matrixplot generated by the gEAR portal (https://umgear.org/p?l=ed724158) showing mean expression values of *Tnfaip2* and *pou4f3* in each cell clusters. Cells were collected from P2 and P7 cochlea and utricle, respectively. *Pou4f3* marks all the HCs.

